# Single Cell Transcriptomics of Pancreatic Cancer Precursors Demonstrates Epithelial and Microenvironmental Heterogeneity as an Early Event in Neoplastic Progression

**DOI:** 10.1101/306134

**Authors:** Vincent Bernard, Alexander Semaan, Jonathan Huang, F. Anthony San Lucas, Feven C Mulu, Bret M Stephens, Paola A. Guerrero, Yanqing Huang, Jun Zhao, Nabiollah Kamyabi, Subrata Sen, Paul A Scheet, Cullen M Taniguchi, Michael P Kim, Ching-Wei Tzeng, Matthew H Katz, Aatur D Singhi, Anirban Maitra, Hector A Alvarez

## Abstract

**Background:** Early detection of pancreatic ductal adenocarcinoma (PDAC) remains elusive. Precursor lesions of PDAC, specifically, intraductal papillary mucinous neoplasms (IPMNs) represent a *bona fide* pathway to invasive neoplasia, although the molecular correlates of progression remain to be fully elucidated. Single cell transcriptomics provides a unique avenue for dissecting both the epithelial and microenvironmental heterogeneity that accompany multistep progression from non-invasive IPMNs to PDAC.

**Methods:** Single cell RNA-sequencing was performed through droplet-based sequencing on 5,403 cells from two low-grade IPMNs (LGD-IPMN), two high-grade IPMNs (HGD-IPMN), and two PDACs (all surgically resected).

**Results:** Analysis of single cell transcriptomes revealed heterogeneous alterations within the epithelium and the tumor microenvironment during the progression of non-invasive dysplasia to invasive cancer. While HGD-IPMNs expressed many core-signaling pathways described in PDAC, LGD-IPMNs harbored subsets of single cells with a transcriptomic profile that overlapped with invasive cancer. Notably, a pro-inflammatory immune component was readily seen in low-grade IPMNs, comprised of cytotoxic T-cells, activated T-helper cells, and dendritic cells, which was progressively depleted during neoplastic progression, accompanied by infiltration of myeloid-derived suppressor cells. Finally, stromal myofibroblast populations were heterogeneous, and acquired a previously described tumor-promoting and immune-evading phenotype during invasive carcinogenesis.

**Conclusions:** This study demonstrates the ability to perform high resolution profiling of the transcriptomic changes that occur during multistep progression of cystic PDAC precursors to cancer. Notably, single cell analysis provides an unparalleled insight into both the epithelial and microenvironmental heterogeneity that accompany early cancer pathogenesis, and might be a useful substrate to identify targets for cancer interception.

## Introduction

Pancreatic ductal adenocarcinoma (PDAC) is the 3rd leading cause of cancer-related deaths in the United States, and most patients present with unresectable disease due to the lack of effective early detection strategies [1]. This reiterates the critical need for understanding the pathogenesis of early neoplasia with the goal of developing biomarkers and molecular targets for cancer interception. The most common cystic neoplasm that is a *bona fide* precursor to PDAC is **I**ntraductal **P**apillary **M**ucinous **N**eoplasms (IPMNs), which comprise roughly 40-50% of resected lesions that are initially diagnosed as asymptomatic pancreatic cysts ^[2]^. While most IPMNs harbor low-grade dysplasia (LGD), it is imperative to distinguish IPMNs that have progressed to high-grade dysplasia (HGD), or harbor an outright invasive component (PDAC). To guide clinicians with identifying IPMNs harboring HGD or PDAC, several radiological “high risk” or “worrisome features” (so-called Sendai and Fukuoka criteria) have been proposed [3]. Although the rate of overdiagnosis and over-treatment have been significantly reduced, these criteria still lack optimal sensitivity and specificity [4]. While patients with non-invasive IPMNs have an excellent prognosis upon surgical resection, once an IPMN develops an invasive component, the probability of long-term survival drops significantly [5]. Although we have now elucidated the signature driver mutations (*KRAS* and *GNAS*) that distinguish IPMNs from other pancreatic cysts, these do not reliably distinguish between indolent *versus* aggressive IPMNs [6, 7]. In fact, the overall state of knowledge remains rudimentary, especially with regards to molecular metrics that can identify ‘aggressive’ precancerous lesions that are likely to progress to carcinoma and require intervention from those that are ‘indolent’ and will naturally regress or remain stable. Thus, much remains to be elucidated in terms of biomarkers of IPMN progression and the underlying molecular features of dysplastic cells that predicate to invasive neoplasia.

The application of single cell DNA and RNA sequencing to cancers has provided unprecedented insights into tumor and microenvironmental heterogeneity present in established cancers. However, the extrapolation of these technologies to the compendium of precursor lesions has been scarce. In the specific context of the pancreas, prior studies have established the transcriptional profiles of normal cell types, such as islet cells, using single cell approaches [8–10]. In this study, we perform the first reported single cell transcriptomic profiling of cystic precursor lesions of PDAC spanning histological grades of dysplastic epithelium. Specifically, we demonstrate our ability to generate transcriptomic libraries from >5400 single cells from surgically resected pancreatic tissues, including two IPMN with LGD, two IPMNs with HGD, and two PDAC lesions utilizing a droplet-based single-cell RNA-seq methodology[11]. Our results demonstrate that epithelial and stromal heterogeneity is evident even within precursor lesions during multistep carcinogenesis, and reflect the progressive co-option of the microenvironment towards a tumor-promoting milieu.

## Results

### Concordance of single cell RNA sequencing to bulk cell lines

Prior to tissue profiling, we investigated the concordance of bulk and single cell RNA sequencing profiles from a PDAC cell line using the droplet-based sequencing technology. Single cell and bulk cell RNA libraries were made using Nextera library preparation chemistry. Single cell RNA-seq was performed at 670 million reads, resulting in 30.4% of the reads mapping to the coding sequence (CDS) regions, and 37.7% mapping to UTR regions with 91,032 reads, 10,800 unique molecular identifier (UMI) counts, and a median of 3,293 unique genes detected per cell passing filter. Correlations in gene expression levels between 2,022 single cells, and bulk cell suspension was excellent (r_s_ > 0.9), with total coverage gene counts of 17,507 and 15,185 for single cells and bulk cells respectively (**Supplementary Figure 1A**). We also verified the concordance of gene counts across two independent replicates of single cell library preparation from the same cell line sample. This revealed a high correlation in gene expression levels (r_s_ > 0.9) with almost identical levels of gene coverage (14,994 and 14,488) (**Supplementary Figure 1B**).

### Preneoplastic epithelium of IPMNs demonstrates both unique and shared transcriptomic signatures with PDAC

We subsequently applied droplet-based single cell RNA sequencing to study the diverse transcriptional profiles that exist within surgically resected preneoplastic (IPMNs) and invasive (PDAC) pancreatic lesions (**Supplementary Figures 1C-D, 2 and Supplementary Table 1**). Cumulatively amongst all of the tissue samples, 5,403 single cells were sequenced. Cells with low expression of genes (<300 genes) and high percentage of mitochondrial genes expressed (>10%) were digitally filtered out resulting in, 3,343 single cells used for the subsequent analysis. Scatter plots of number of UMIs compared to number of genes and abundance of mitochondrial transcripts revealed consistent read depth across single cells between lesions and absence of apoptosis induced transcript batch effects (**Supplementary Figure 1C**). The mean number of genes and UMIs detected per cell was 1101 and 3194 respectively. After identifying the top variable genes, we performed principal component analysis (PCA) and determined which principal components (**Supplementary Figure 1D**) to use for unsupervised clustering using t-distribution stochastic neighbor embedding (t-SNE)[12] which was implemented using the SEURAT package (**Figure 1A**) [11]. This analysis identified ten distinct subpopulations (“clusters”) comprised of unique stromal and epithelial components classified defined by characteristic gene expression patterns (**Figure 1B-C**).

**Figure 1:**
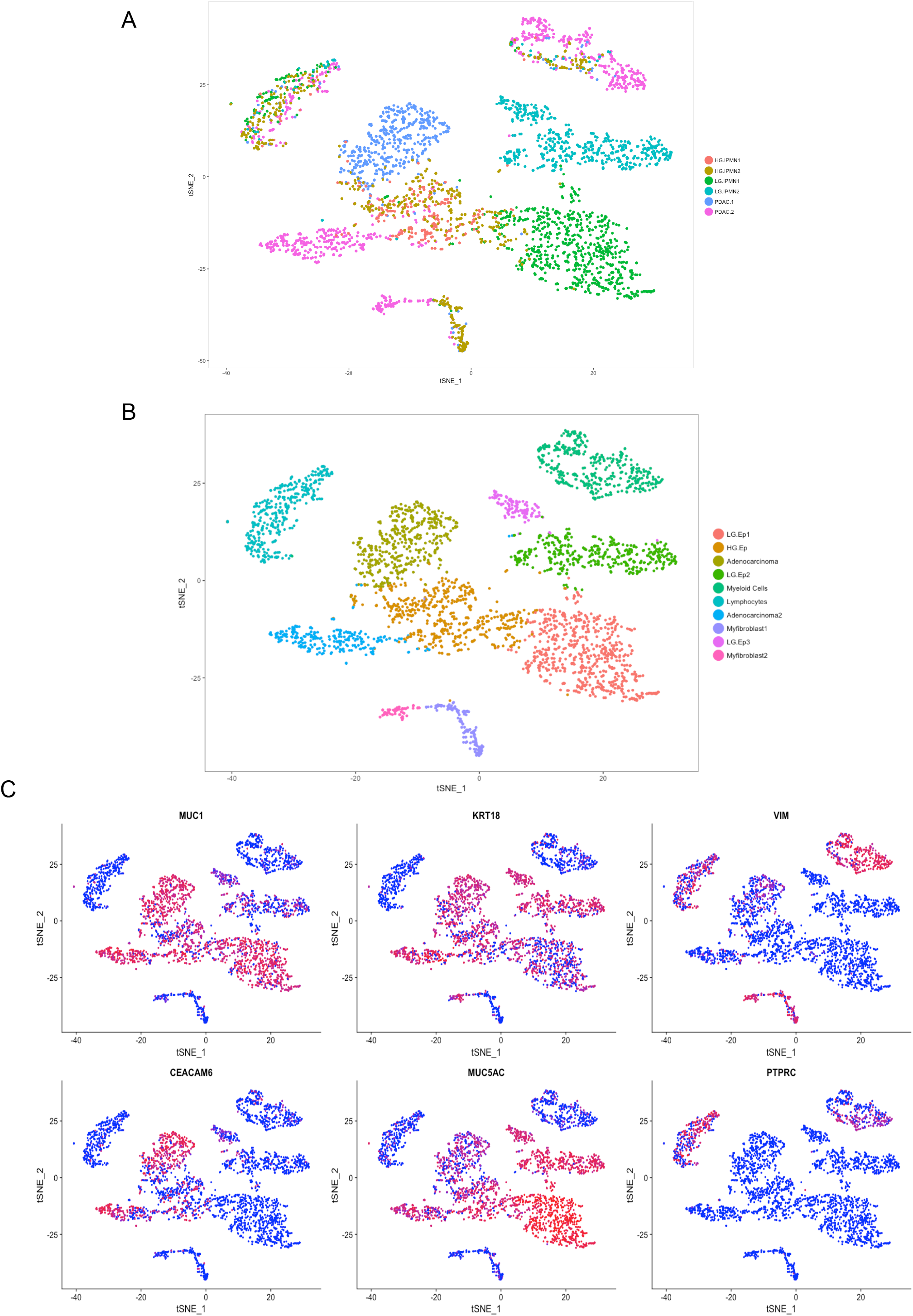
tSNE plots of all 3,343 cells from six lesions included in this study, **(A)** colors represent different tissue samples **(B)** colors represent unique cell types characterized by gene expression (Ductal Epithelium = Ep). **(C)** Feature plots demonstrating expression of specified genes among clusters on the tSNE.

We first identified the neoplastic single cell “clusters” through previously defined signature transcripts for pancreatic epithelial lesions, including *KRT19* and *MUC1*, which were present among all samples types (IPMNs and PDAC) irrespective of grade if dysplasia or invasion[13, 14] (**Figure 2A**). We subsequently sought to correlate the histological grading of the designated “clusters” with known biomarkers of dysplastic progression to cancer. This revealed high expression of transcripts like *CEACAM6* within subpopulations of HGD-IPMN and PDAC samples, which confirmed previously published data on this marker in bulk RNA analysis and immunohistochemistry of intact tissues [15, 16]. Conversely, we observed high expression of *MUC5AC*, which encodes for an apomucin mostly seen in LGD and downregulated during histological progression, within the LGD-IPMNs compared to the other sample types [13].

**Figure 2:**
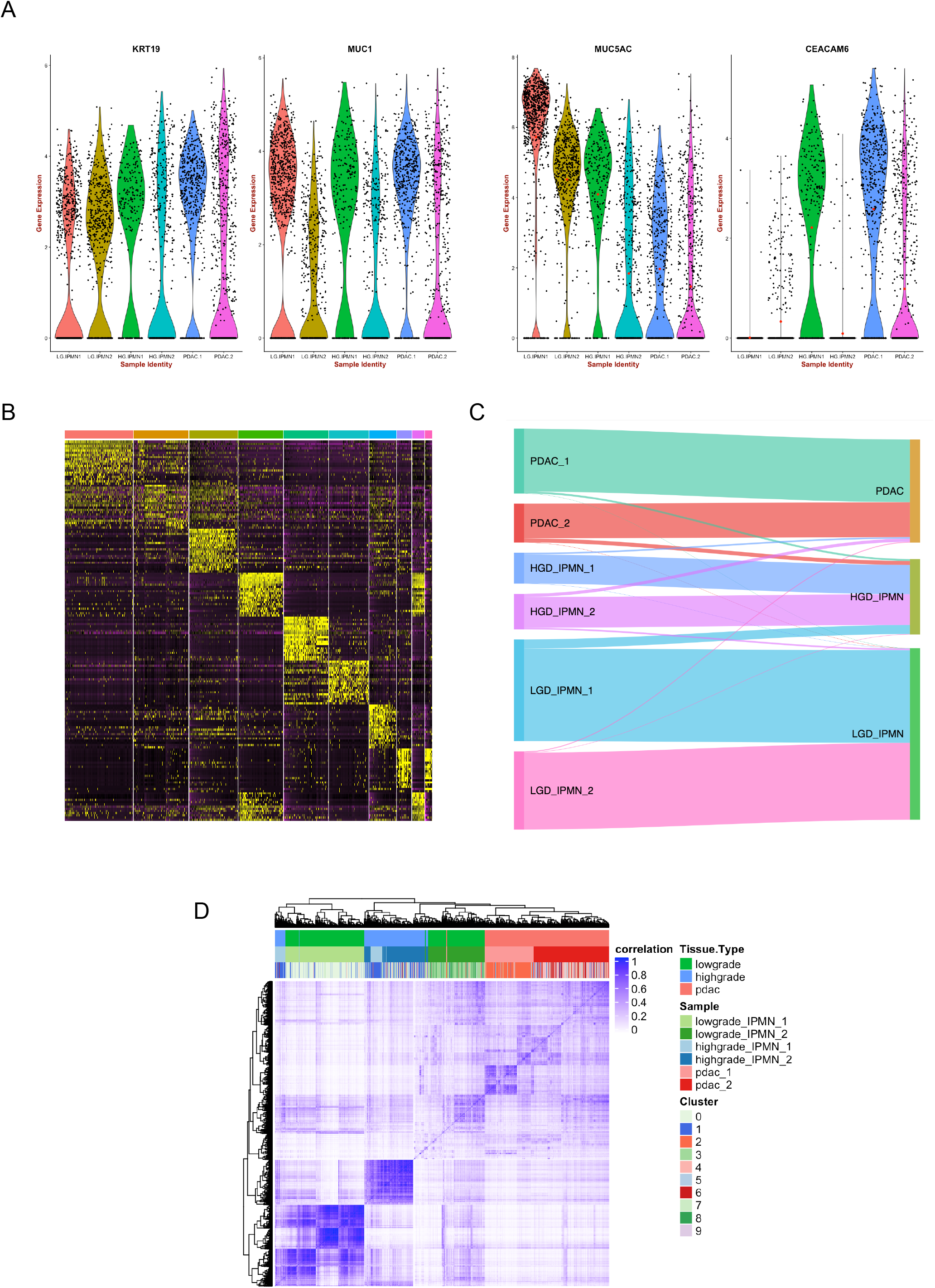
**(A)** Violin plots confirm expression of characteristic PDAC and cystic preneoplasia related markers across lesion types. **(B)** Heatmap of the top 25 differentially expressed genes used to identify cell phenotypes across ten discrete clusters. **(C)** Sankey diagram demonstrating epithelial cells profiled from LGD-IPMNs, HGD-IPMNS, and PDAC tissue and where they reside within annotated tSNE clusters **(D)** Correlation heatmap individual cells across all lesions, identified by originating lesion type and tSNE cluster.

In an effort to identify “cluster-defining” signatures, we profiled the top 25 differentially expressed genes within each of the ten distinct single cell “cluster” (**Figure 2B, Supplementary Table 2**). Annotation of the resulting transcripts revealed aberrant expression of multiple cancer-related genes even within the LGD-IPMN cells (designated as “clusters” LG.Ep1, 2, and 3 (**Figure 1B**). These include overexpression of transcripts such as *TFF3 and REG4* that have been previously described as upregulated during cancer progression[17, 18]. On the contrary, the LGD-IPMN clusters demonstrated persistent expression of putative tumor suppressor genes (TSGs) such as *RAP1GAP* that have been shown to suppress tumor invasion and metastases across various cancers (including RAS mutant neoplasms) [19], while HGD-IPMNs (HG.Ep) revealed downregulation of the aforementioned TSGs, concomitant with higher expression of oncogenic transcripts previously associated with progression to PDAC such as *S100P* and *S100A10*, amongst others [20–22].

We then investigated whether biological differences could be observed through gene ontology and pathway analysis (IPA: Ingenuity Pathways Analysis) of differentially expressed genes between neoplastic epithelial clusters (**Figure 1B**). Several aberrant canonical pathways were identified during the transition from LGD to HGD IPMN that are related to previously published core signaling pathways in PDAC, including integrin signaling, signaling by small GTPases, Wnt/B-catenin signaling, axonal guidance signaling, apoptosis, and G1/S phase regulation (**Supplementary Table 3**) [23, 24]. When comparing the PDAC and LGD lesions, the differential expression between all of these pathways was maintained, with the additional deregulation of DNA damage response, TGF-B signaling, and SAPK/JNK signaling. Other aberrant canonical pathways identified during transition from LGD to HGD and PDAC lesions include metabolism related pathways such as oxidative phosphorylation and mitochondrial dysfunction, as well as mTOR signaling (**Supplementary Table 4**).

It is important to note that even “adenocarcinoma”-designated clusters contained cells originating from LGD-IPMNs with associated expression of PDAC core signaling pathways and deregulation of tumor suppressor genes (**Figure 2C**). In our particular data set, 1.2% of epithelial cells from LGD-IPMNs were found in adenocarcinoma related clusters. Additionally, up to 8.9% of cells from LGD-IPMNs tended to cluster with HGD-IPMNs with their respective changes in gene expression pathways. This suggests that even within IPMNs with otherwise LGD histology, there are cells that phenocopy the transcriptomic features of invasive neoplasia (**Supplementary Figure 2**). The expression profiles of such low frequency cells within LGD-IPMNs would likely be missed during bulk RNA sequencing, and further underscores the utility of the single cell sequencing approach in elucidating the epithelial heterogeneity that exists even within early precursor lesions of PDAC.

Analysis of the cell-to-cell correlations for gene expression of the 3,343 cells demonstrated relatively higher intra-tumoral coherence among cells from LGD and HGD lesions compared with those from PDAC (**Figure 2D**), a not surprising finding suggesting an increase in intra-tumoral epithelial heterogeneity during the progression from IPMNs to PDAC [25]. Inter-lesion correlation was better observed in cells derived from stromal components including myeloid and lymphocytic populations, whereby a significant number of populations showed similarities across tissue types (**Figure 1A-C**). This suggests the presence of common cancer associated immune components among lesion types. On the other hand, even though these stromal components tended to cluster with one another (**Figure 1B**), correlation heat maps suggested the presence of multiple unique subtypes within the stroma and non-random variations during histological progression to cancer (**Supplementary Figure 4**, *and results below*).

### Virtual microdissection of stromal and immune heterogeneity during IPMN progression

In an effort to better identify unique subpopulations of stromal components across lesion types, we opted to perform single cell digital microdissection of only stromal cells and exclude epithelial components. This resulted in the identification of seven unique clusters with varying degrees of enrichment across lesions types (**Figures 3a,b**). Differential expression of the top 25 genes across clusters allowed us to identify distinct immune and myofibrolast-derived phenotypes within each lesional subtype (**Figure 3c,d**).

**Figure 3:**
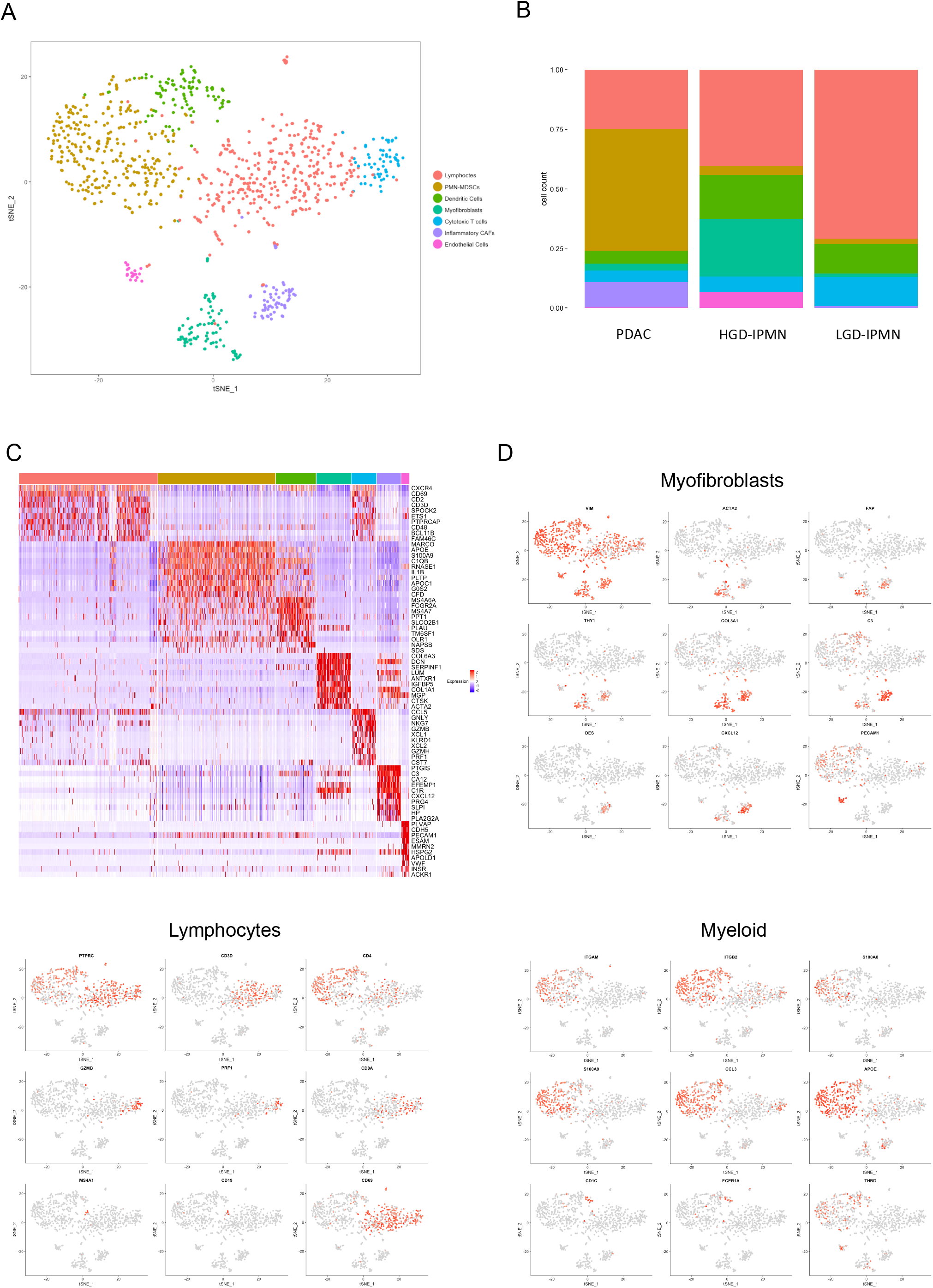
**(A)** tSNE plot of all stromal cells that were virtually microdissection from entire lesions. Different colors represent unique cell phenotypes. **(B)** Proportion of cell phenotypes enriched in each lesion (PDAC, HG IPMN, and LG IPMN) **(C)** Heatmap of the top 20 differentially expressed genes used to identify cell phenotypes across clusters. **(D)** Feature Plots demonstrating expression of specific genes among clusters to identify respective cell types.

A high proportion of cytotoxic T cells (measured by CD8, and presence of granzyme and perforin related transcripts) were observed in LGD-IPMNs compared to HGD-IPMNs and PDAC. In proportion to other immune subtypes, CD4 T cells also appear to be more highly enriched in LGD-IPMN compared to others, and present with generalized activation as defined by expression of *CD69*. We also detected the presence of rare B-cell populations (expressing CD20 and CD19) that are present in both HGD-IPMNs and in LGD-IPMNs, but are completely absent in PDACs. Presence of tumor infiltrating B-cells have recently been described in PanIN lesions with an immunosuppressive role during the initiating stages of PDAC multistep progression, and may, in fact, have a similar role in the context of IPMNs based on these findings [26–28].

Notably, we observed a significantly enriched proportion of myeloid derived suppressor cells (MDSCs), within the stromal component of PDAC, representing 51% (277/544) of single stromal cells profiled, compared to 2.3% (3/131) and 3.5% (10/281) within LGD-IPMNs and HGD-IPMNs respectively. These MDSCs resemble the pro-tumorigenic polymorphonuclear myeloid-derived suppressor cell (PMN-MDSC) phenotype based on expression of CD11b (*ITGAM*), S100A9, CCL3, and APOE, which has been previously described to be prevalent during cancer progression [29]. Among myeloid derived populations, we also observed cDC2 type dendritic cells, characterized by expression of CD1C, THBD, and *FCER1A* [30]. These cells have been shown to have T-cell stimulatory potential and are critical mediators of cross presentation of tumor antigens mediated through the high-affinity IgE receptor FcεRI (*FCER1A*) [31]. This pro-inflammatory subpopulation appears in greater proportions amongst LGD and HGD-IPMNs, suggesting a more prevalent predication for anti-tumor immune response within pre-neoplastic lesions.

Heterogeneous fibroblast populations across histological subtypes were also identified potentially representing distinct stromal functions during tumorigenesis. For example, the fibroblast subtype known as "inflammatory” CAF (iCAF) is characterized by expression of *VIM, FAP, COL3A1, DES, IL6*, and *CXCL12* and reduced expression of a-SMA (*ACTA2*)[32, 33]. This subpopulation has been shown to be involved in immunosuppression, growth factor secretion, and promotion of pro-tumorigenic mechanisms facilitating invasion and metastasis[32], which might correlate with its exclusive representation in PDAC derived clusters, representing 10.5% (58/544) of single stromal cells profiled, and absence within non-invasive IPMNs. A separate compartment of CAFs, described as myofibroblasts (“myCAFs”), with increased alpha-SMA expression and reduced expression of *CXCL12* and *DES*, was also identified. These myCAFs have been implicated in distinct functions from iCAFs, including secretion of autocrine stromal and endothelial growth factors [32]. This population is rare in LGD-IPMNs, but is highly represented in HGD-IPMNs suggesting that activation of fibroblasts to the myCAF phenotype tends to occur even within non-invasive dysplastic lesions (**Figure 4**).

**Figure 4:**
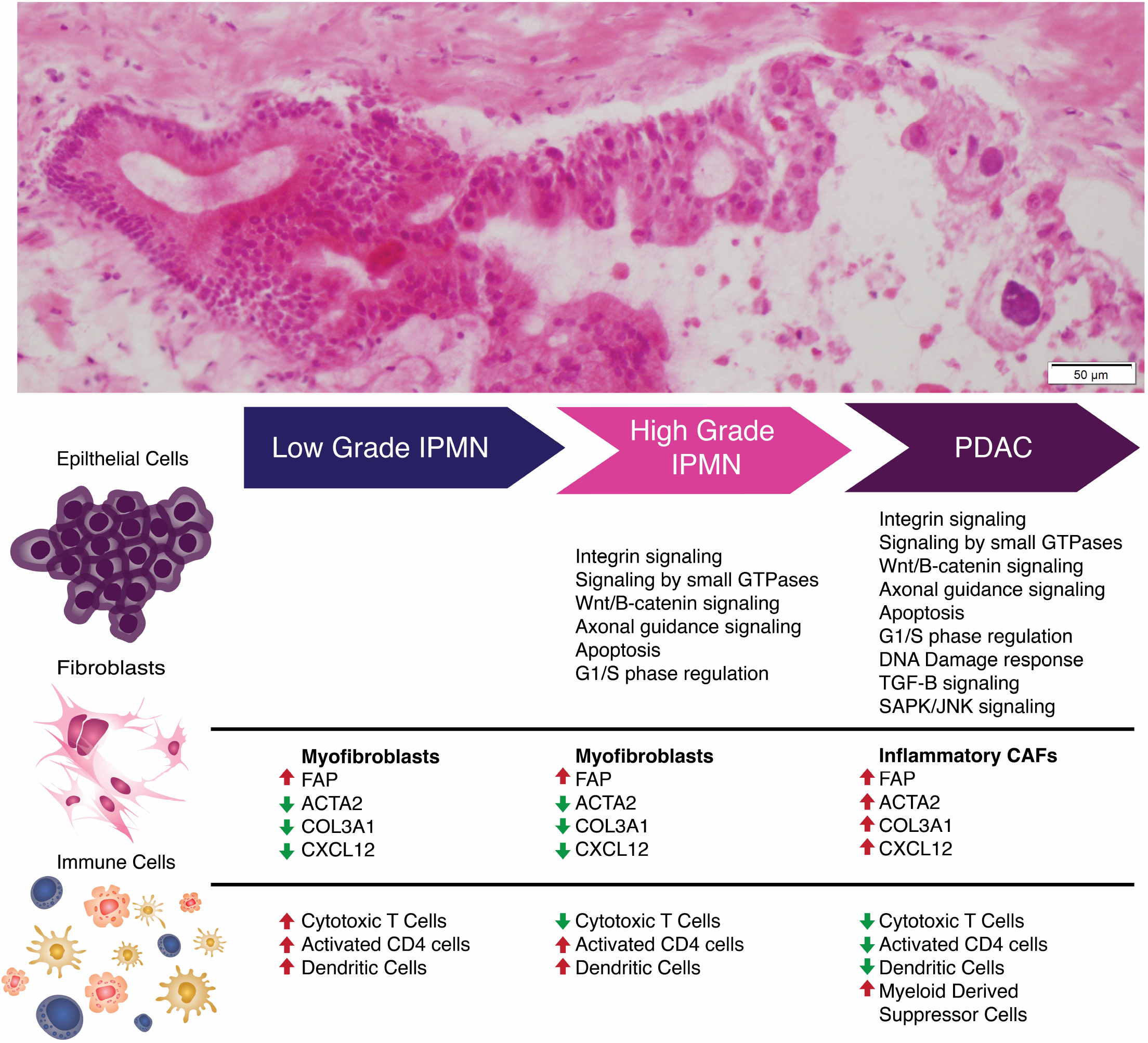
Schematic representation of evolving molecular and phenotypic signatures during preneoplastic progression of pancreatic cancer.

## Discussion

Histologically well-established precancerous lesions precede the onset of PDAC by years, if not decades. Although much has been described in regards to the genomic alterations in these precursor lesions, the overall state of knowledge remains rudimentary, especially with regards to molecular metrics that can identify ‘aggressive’ precancerous lesions that require intervention from those that are ‘indolent’ and will naturally regress or remain stable. In the current study, we describe an approach at profiling the molecular events that occur during pancreatic carcinogenesis in the context of cystic precursors (IPMNs) with the goal of understanding the heterogeneity of epithelial and stromal components that might delineate lesions with an aggressive potential. We do this through high-resolution single cell RNA sequencing of LGD-IPMN, HGD-IPMN, and PDACs, which allows us to specifically profile aberrant pathways across multiple cell types. Interestingly, progression from LGD and HGD lesions to invasive PDAC revealed shifts in distinct cell populations encompassing both epithelial and stromal/immune compartments.

Among epithelial populations, we detected expression of transcripts of gastric lineage that have been previously described during IPMN progression and have been correlated with better prognosis, such as *MUC5AC*, as differentially overexpressed in our LGD lesions [34]. Although oncogenic transcripts are also expressed at the LGD stage, there appears to be retained expression of tumor suppressor related pathways which may help counteract further dysplastic progression of these lesions. Within HGD lesions, differential expression of these tumor suppressor pathways are no longer detected and we begin to find previously described classical core signaling pathways in PDAC [24, 35].

As a stroma-rich cancer, multiple studies have shown how the diverse tumor microenvironment (TME) plays a crucial role in the development, progression, and immune evasion that exemplifies the biology of PDAC [36, 37]. As previously shown by Moffit et al., “virtual microdissection” of stromal genes revealed an “activated” subpopulation of stroma characterized by overexpression of inflammatory pathways [38]. Although these findings are further validated within our own study, Moffit et al., were not able to distinguish specific contributions of immune and myofibroblast components to this signature. By dissecting specific cell-types at the single cell level, we further delineate this signature to describe the dichotomy of “myCAF” *versus* “iCAF” fibroblast populations during multistep progression, with the former cells observed (albeit rarely) even in LGD-IPMNs and the latter cells only identified in PDAC samples. Within our analysis, besides expression of many typical CAF markers (*FAP, THY1*, and *DES*), the iCAF cells also show a unique expression pattern of the C-X-C motif chemokine ligand 12 (CXCL12) [39]. This chemokine and its corresponding receptor (CXCR4) are believed to play a crucial role in many solid cancers, and are associated with metastatic spread in breast cancer, lung cancer and melanoma [40–43]. Accumulating evidence also points to a role in PDAC progression, with higher expression of CXCL12 correlating with metastasis, likely by facilitating immune evasion, and increased levels of matrix metalloproteinases leading to cellular invasion [44–47]. Within our own study, we find similar trends in the presence of other cellular populations that parallel the emergence of CXCL12-expressing iCAFs, such as a decrease in cytotoxic T cell and increase in myeloid suppressive proportions, creating the notorious immune suppressive TME well described in PDAC.

Myeloid derived subpopulations also tend to evolve during tumor progression. Comprised of multiple clusters expressing CD11b, *ITGB2*, *CD13*, *CD18*, and *S100A9*, a clear separation based on their individual transcriptomes remains difficult. This high overlap of myeloid derived subpopulations mirrors recent findings in breast cancer where previously proposed M1 and M2 macrophage associated genes are frequently expressed within the same cells among these two clusters [48]. As Azizi et al, describes, this suggests that distinct prototypical macrophages may not be prevalent within tumors and may in fact exist within the spectrum of these two phenotypes. Among other myeloid derived populations, we do identify PMN-MDSCs which are known surrogates of prognosis and contribute to immune evasion and tumor progression through neo-angiogenesis, migration and invasion, and thus metastatic spread [49, 50]. Similar to the elevated proportion of CAFs experienced in more advanced disease we describe a striking increase in MDSC populations compared to pre-neoplastic lesions (51% of non-epithelial cell types in PDAC, versus <5% in non-invasive IPMNs). This resembles findings from Kumar et al. whereby CAFs are able to actively recruit PMN-MDSC to tumors, further supporting the role of an “inflammatory CAF” subpopulation in promoting immunosuppression [51]. The identification of demonstrable dendritic cell (DC) populations within LGD and HGD IPMN microenvironment suggests a pro-inflammatory phase that is present during the non-invasive precursor states. This is particularly true in LGD IPMNs where a higher proportion of cytotoxic T cells exist in comparison to other lesion types that may be supported by cDC2 cells through cross presentation of tumor antigens and T-cell stimulation. Additionally, the presence of these dendritic cells in IPMNs may also provide opportunities for cancer “immune interception”, since dendritic cell targeted vaccines have previously shown effectiveness in the context of an myeloid immunosuppressive environment through reduction of PMN-MDSCs [52].

In conclusion, we describe how the TME may evolve during the multistep progression of IPMNs to PDAC, whereby the non-invasive precursor lesions begin to experience a loss of cytotoxic T-cells during dysplastic progression and begin to assimilate immune components such as PMN-MDSCs, with immune suppressive properties. Additionally, we demonstrate this permissive microenvironment is correlated with appearance of tumor-promoting iCAF stromal cell populations that facilitate immune evasion. Notably, single cell analysis of IPMNs reveals, for the first time, that even in otherwise histologically innocuous LGD-IPMNs, we find minor populations of cells that transcriptionally phenocopy HGD and PDAC (~9% and ~1% of LGD-IPMN cells, respectively). It is possible that future single cell analyses on a larger series of LGD-IPMNs might establish a “threshold” which portends aggressive natural history even in the absence of radiologically detectable worrisome features. Although it is important to stress the limited generalizations that can be made from such a small subset of lesions, the ability to detect these gene expression patterns among single cells provides a primer for uncovering how heterogeneous cell types contribute to tumor carcinogenesis. Leveraging this strategy may thus facilitate elucidation of molecular biomarkers for disease stratification.

## Materials and methods

### Preparation of fresh tissue material from surgically resected IPMNs or PDAC

A total of six patients were recruited at MD Anderson Cancer Center (MDACC) and University of Pittsburgh Medical Center (UPMC) through informed consent following institutional review board (IRB) approval at both institutions (PA15-0014, Lab08-0098, Lab05-0080, Lab00-396). Two PDACs and one HGD-IPMN was profiled from MDACC and two LGD-IPMNs and one HGD-IPMN was profiled from UPMC. Pancreatic tissue was delivered to the laboratory after surgical resection in DMEM, high glucose, GlutaMAX™ Supplement, HEPES (Thermo fisher, 10564011) in 1% Bovine Serum Albumin (Thermo fisher, B14) in a 15ml conical tube. Tissue was then transferred to a 35×12mm Petri Dish(Thermo fisher, #150318), and minced with sterile surgical scalpel to 0.5-1.0mm fragments in approximately 1ml of the media.

Tissue digestion was performed with Liberase™ TH Research Grade (Sigma-Aldrich, 5401135001) alone for IPMN samples, or with both Liberase™ TH Research Grade and Accutase^®^ solution for PDAC tissues (Sigma-Aldrich, A6964). For warm digestion with Liberase™ TH Research Grade, pancreatic tissue fragments were incubated to a final concentration of 10 mg/ml, and placed on a shaker at 37 °C, 225 RPM for 20 min and gently pipetted every 10 min. At the end of the digestion period, the fragments (tissue slurry) were gently pipetted and washed to maximize the release of single cells. The tissue slurry was passed through a 100um cell strainer followed by a 35um cell strainer. The single-cell suspension was transferred to a new tube and centrifuged for 5 min at 1500 RPMI at 4°C. The supernatant was discarded and the cell pellet was re-suspended in 400 ul of PBS for downstream cell viability analysis and cell counting.

For warm digestion of PDAC tissues, we followed the identical procedure above as the IPMNs, followed by a second digestion period using sterile-filtered Accutase^®^ solution, and placed on a shaker at 37 °C, 225 RPM for 30min, with gentle pipetting every 10 minutes.

### ddSEQ Surecell Library Preparation

Single cell transcriptomic amplification and library prep was performed using the SureCell WTA 3’ Library Prep Kit for the ddSEQ System (Illumina, Cat# 20014279). Briefly, single cells were individually partitioned into sub-nanoliter droplets. Single cells were then lysed and barcoded inside individual droplets, with subsequent first strand synthesis. Individual droplets containing barcoded sample cells were disintegrated, purified, and subject to second strand cDNA synthesis. Purified amplified cDNA was subsequently quantified on a 2200 TapeStation System (Agilent) in order to validate quality of amplification. Successful reactions were then fragmented and amplified using Nextera technology for low input cDNA material. Libraries were purified, quantified, and then sequenced on the Illumina NextSeq 500.

### Single cell clustering and transcriptomic analyses

Resulting FastQ files were run through the Single cell RNA-seq app on BaseSpace. Briefly, reads were aligned against a reference genome using Spliced Transcripts Alignment to a Reference (STAR), followed by barcode tagging and BAM indexing [53]. Count files are generated using gene UMI counter and with cells passing quality filter based on cells above background and passing knee filter. Gene ontology and KEGG pathway analyses are performed using annotation functions of the R limma package.

To perform the T-SNE clustering and additional downstream analyses, the UMI count files were compiled into sparse matrices and subsequently filtered based on the criteria that each cell must express a minimum of 300 genes and be composed of less than 10% mitochondrial genes. The data was then log normalized, followed by the identification of variable genes. The number of UMIs and the percentage of mitochondrial genes was regressed out, and the resulting data was scaled and centered. The sparse matrix is analyzed by PCA and then run through a jackstraw algorithm to estimate statistical significance between the genes and principal components. After using this analysis to determine the number of dimensions to use, the result was clustered in a shared nearest neighbor algorithm and T-SNE before plotting. These annotated matrices were also used to find differentially expressed genes between subpopulations. The resulting T-SNE plots were colored according to specific features to visually present expression of genes in various clusters. Functions for analysis provided in the Seurat package [11].

The correlation matrices were created based on the filtered list of cells used in the T-SNE. Matrices were formed through the function 'cor' from the stats package. The Pearson correlation coefficients were then plotted in ComplexHeatmap[54]. Cells are ordered through hierarchical clustering. Both T-SNE and correlation heatmaps were created in R v3.4.2

**Supplementary Figure 1: (A)** Correlation of gene expression levels in a pancreatic cancer cell line between single cells, bulk cells, and across independent replicates. **(B)** Scatter plots of percentage mitochondria, number of UMIs, and number of genes expressed per single cell across independent tissue samples from pancreatic lesions (different colors). **(C)** Violin plots of number of genes, UMIs, and percentage of mitochondrial genes expressed per single cell from tissue samples profiled in this study. **(D)** Principal components (PC) elbow plot to determine the elbow between the standard deviation of the PC and the number of PCs (11).

**Supplementary Figure 2 (A)** Gross pathology of a HGD-IPMN lesion with concomitant PDAC included in this study. **(B)** Paraffin fixed tissue H&E sections of representative LGD-IPMN and HGD-IPMN lesions included in this study (20x magnification).

**Supplementary Figure 3:** Feature Plot of characteristic genes plotted across tSNE clusters demonstrating stromal phenotypes.

**Supplementary Figure 4:** Correlation heatmap of individual cells across stromal populations identified by originating lesion type and tSNE cluster.

## Acknowledgements

This work was supported by the MD Anderson Moonshot Program and the Khalifa Bin Zayed Al-Nahyan Foundation; the National Institute of Health (U01CA196403 and U01CA200468 to AM); the Cancer Prevention Research Institute of Texas (RP140106 to VB), the DFG (German Research Foundation) (SE-2616/2-1 to A.S).

## References

1. Siegel RL, Miller KD, Jemal A: Cancer statistics, 2018. CA Cancer J Clin 2018.

2. Salvia R, Crippa S, Partelli S, Armatura G, Malleo G, Paini M, Pea A, Bassi C: Differences between main-duct and branch-duct intraductal papillary mucinous neoplasms of the pancreas. World J Gastrointest Surg 2010, 2(10):342–346.

3. Tanaka M, Fernandez-Del Castillo C, Kamisawa T, Jang JY, Levy P, Ohtsuka T, Salvia R, Shimizu Y, Tada M, Wolfgang CL: Revisions of international consensus Fukuoka guidelines for the management of IPMN of the pancreas. Pancreatology 2017, 17(5):738–753.

4. Heckler M, Michalski CW, Schaefle S, Kaiser J, Buchler MW, Hackert T: The Sendai and Fukuoka consensus criteria for the management of branch duct IPMN - A meta-analysis on their accuracy. Pancreatology 2017, 17(2):255–262.

5. Rezaee N, Barbon C, Zaki A, He J, Salman B, Hruban RH, Cameron JL, Herman JM, Ahuja N, Lennon AM et al: Intraductal papillary mucinous neoplasm (IPMN) with high-grade dysplasia is a risk factor for the subsequent development of pancreatic ductal adenocarcinoma. HPB (Oxford) 2016, 18(3):236–246.

6. Molin MD, Matthaei H, Wu J, Blackford A, Debeljak M, Rezaee N, Wolfgang CL, Butturini G, Salvia R, Bassi C et al: Clinicopathological correlates of activating GNAS mutations in intraductal papillary mucinous neoplasm (IPMN) of the pancreas. Ann Surg Oncol 2013, 20(12):3802–3808.

7. Matthaei H, Wu J, Dal Molin M, Shi C, Perner S, Kristiansen G, Lingohr P, Kalff JC, Wolfgang CL, Kinzler KW et al: GNAS sequencing identifies IPMN-specific mutations in a subgroup of diminutive pancreatic cysts referred to as “incipient IPMNs”. Am J Surg Pathol 2014, 38(3):360–363.

8. Segerstolpe A, Palasantza A, Eliasson P, Andersson EM, Andreasson AC, Sun X, Picelli S, Sabirsh A, Clausen M, Bjursell MK et al: Single-Cell Transcriptome Profiling of Human Pancreatic Islets in Health and Type 2 Diabetes. Cell Metab 2016, 24(4):593–607.

9. Muraro MJ, Dharmadhikari G, Grun D, Groen N, Dielen T, Jansen E, van Gurp L, Engelse MA, Carlotti F, de Koning EJ et al: A Single-Cell Transcriptome Atlas of the Human Pancreas. Cell Syst 2016, 3(4):385–394 e383.

10. Li J, Klughammer J, Farlik M, Penz T, Spittler A, Barbieux C, Berishvili E, Bock C, Kubicek S: Single-cell transcriptomes reveal characteristic features of human pancreatic islet cell types. EMBO Rep 2016, 17(2):178–187.

11. Macosko EZ, Basu A, Satija R, Nemesh J, Shekhar K, Goldman M, Tirosh I, Bialas AR, Kamitaki N, Martersteck EM et al: Highly Parallel Genome-wide Expression Profiling of Individual Cells Using Nanoliter Droplets. Cell 2015, 161(5):1202–1214.

12. Waltman LvE, N. J: A smart local moving algorithm for large-scale modularity-based community detection. Eur Phys 2013, 86, 471

13. Jonckheere N, Skrypek N, Van Seuningen I: Mucins and pancreatic cancer. Cancers (Basel) 2010, 2(4):1794–1812.

14. Zhang T, Wang X, Yue Z: Identification of candidate genes related to pancreatic cancer based on analysis of gene co-expression and protein-protein interaction network. Oncotarget 2017, 8(41):71105–71116.

15. Duxbury MS, Matros E, Clancy T, Bailey G, Doff M, Zinner MJ, Ashley SW, Maitra A, Redston M, Whang EE: CEACAM6 is a novel biomarker in pancreatic adenocarcinoma and PanIN lesions. Ann Surg 2005, 241(3):491–496.

16. Duxbury MS, Ito H, Benoit E, Ashley SW, Whang EE: CEACAM6 is a determinant of pancreatic adenocarcinoma cellular invasiveness. Br J Cancer 2004, 91(7):1384–1390.

17. Terris B, Blaveri E, Crnogorac-Jurcevic T, Jones M, Missiaglia E, Ruszniewski P, Sauvanet A, Lemoine NR: Characterization of gene expression profiles in intraductal papillary-mucinous tumors of the pancreas. Am J Pathol 2002, 160(5):1745–1754.

18. He XJ, Jiang XT, Ma YY, Xia YJ, Wang HJ, Guan TP, Shao QS, Tao HQ: REG4 contributes to the invasiveness of pancreatic cancer by upregulating MMP-7 and MMP-9. Cancer Sci 2012, 103(12):2082–2091.

19. Zhang L, Chenwei L, Mahmood R, van Golen K, Greenson J, Li G, D'Silva NJ, Li X, Burant CF, Logsdon CD et al: Identification of a putative tumor suppressor gene Rap1GAP in pancreatic cancer. Cancer Res 2006, 66(2):898–906.

20. Arumugam T, Simeone DM, Van Golen K, Logsdon CD: S100P promotes pancreatic cancer growth, survival, and invasion. Clin Cancer Res 2005, 11(15):5356–5364.

21. Kim GE, Bae HI, Park HU, Kuan SF, Crawley SC, Ho JJ, Kim YS: Aberrant expression of MUC5AC and MUC6 gastric mucins and sialyl Tn antigen in intraepithelial neoplasms of the pancreas. Gastroenterology 2002, 123(4):1052–1060.

22. Sitek B, Sipos B, Alkatout I, Poschmann G, Stephan C, Schulenborg T, Marcus K, Luttges J, Dittert DD, Baretton G et al: Analysis of the pancreatic tumor progression by a quantitative proteomic approach and immunhistochemical validation. J Proteome Res 2009, 8(4):1647–1656.

23. Bailey P, Chang DK, Nones K, Johns AL, Patch AM, Gingras MC, Miller DK, Christ AN, Bruxner TJ, Quinn MC et al: Genomic analyses identify molecular subtypes of pancreatic cancer. Nature 2016, 531(7592):47–52.

24. Biankin AV, Waddell N, Kassahn KS, Gingras MC, Muthuswamy LB, Johns AL, Miller DK, Wilson PJ, Patch AM, Wu J et al: Pancreatic cancer genomes reveal aberrations in axon guidance pathway genes. Nature 2012, 491(7424):399–405.

25. Oldfield LE, Connor AA, Gallinger S: Molecular Events in the Natural History of Pancreatic Cancer. Trends Cancer 2017, 3(5):336–346.

26. Pylayeva-Gupta Y, Das S, Handler JS, Hajdu CH, Coffre M, Koralov SB, Bar-Sagi D: IL35-Producing B Cells Promote the Development of Pancreatic Neoplasia. Cancer Discov 2016, 6(3):247–255.

27. Lee KE, Spata M, Bayne LJ, Buza EL, Durham AC, Allman D, Vonderheide RH, Simon MC: Hif1a Deletion Reveals Pro-Neoplastic Function of B Cells in Pancreatic Neoplasia. Cancer Discov 2016, 6(3):256–269.

28. Gunderson AJ, Kaneda MM, Tsujikawa T, Nguyen AV, Affara NI, Ruffell B, Gorjestani S, Liudahl SM, Truitt M, Olson P et al: Bruton Tyrosine Kinase-Dependent Immune Cell Cross-talk Drives Pancreas Cancer. Cancer Discov 2016, 6(3):270–285.

29. Ouzounova M, Lee E, Piranlioglu R, El Andaloussi A, Kolhe R, Demirci MF, Marasco D, Asm I, Chadli A, Hassan KA et al: Monocytic and granulocytic myeloid derived suppressor cells differentially regulate spatiotemporal tumour plasticity during metastatic cascade. Nat Commun 2017, 8:14979.

30. Villani AC, Satija R, Reynolds G, Sarkizova S, Shekhar K, Fletcher J, Griesbeck M, Butler A, Zheng S, Lazo S et al: Single-cell RNA-seq reveals new types of human blood dendritic cells, monocytes, and progenitors. Science 2017, 356(6335).

31. Platzer B, Elpek KG, Cremasco V, Baker K, Stout MM, Schultz C, Dehlink E, Shade KT, Anthony RM, Blumberg RS et al: IgE/FcepsilonRI-Mediated Antigen Cross-Presentation by Dendritic Cells Enhances Anti-Tumor Immune Responses. Cell Rep 2015.

32. Ohlund D, Handly-Santana A, Biffi G, Elyada E, Almeida AS, Ponz-Sarvise M, Corbo V, Oni TE, Hearn SA, Lee EJ et al: Distinct populations of inflammatory fibroblasts and myofibroblasts in pancreatic cancer. J Exp Med 2017, 214(3):579–596.

33. Avery D, Govindaraju P, Jacob M, Todd L, Monslow J, Pure E: Extracellular matrix directs phenotypic heterogeneity of activated fibroblasts. Matrix Biol 2017.

34. Yonezawa S, Horinouchi M, Osako M, Kubo M, Takao S, Arimura Y, Nagata K, Tanaka S, Sakoda K, Aikou T et al: Gene expression of gastric type mucin (MUC5AC) in pancreatic tumors: its relationship with the biological behavior of the tumor. Pathol Int 1999, 49(1):45–54.

35. Jones S, Zhang X, Parsons DW, Lin JC, Leary RJ, Angenendt P, Mankoo P, Carter H, Kamiyama H, Jimeno A et al: Core signaling pathways in human pancreatic cancers revealed by global genomic analyses. Science 2008, 321(5897):1801–1806.

36. Kalluri R: The biology and function of fibroblasts in cancer. Nat Rev Cancer 2016, 16(9):582–598.

37. Chang JH, Jiang Y, Pillarisetty VG: Role of immune cells in pancreatic cancer from bench to clinical application: An updated review. Medicine (Baltimore) 2016, 95(49):e5541.

38. Moffitt RA, Marayati R, Flate EL, Volmar KE, Loeza SG, Hoadley KA, Rashid NU, Williams LA, Eaton SC, Chung AH et al: Virtual microdissection identifies distinct tumor- and stroma-specific subtypes of pancreatic ductal adenocarcinoma. Nat Genet 2015, 47(10):1168–1178.

39. Shiga K, Hara M, Nagasaki T, Sato T, Takahashi H, Takeyama H: Cancer-Associated Fibroblasts: Their Characteristics and Their Roles in Tumor Growth. Cancers (Basel) 2015, 7(4):2443–2458.

40. Xu C, Zhao H, Chen H, Yao Q: CXCR4 in breast cancer: oncogenic role and therapeutic targeting. Drug Des Devel Ther 2015, 9:4953–4964.

41. Jiang YM, Li G, Sun BC, Zhao XL, Zhou ZK: Study on the relationship between CXCR4 expression and perineural invasion in pancreatic cancer. Asian Pac J Cancer Prev 2014, 15(12):4893–4896.

42. Tsai MF, Chang TH, Wu SG, Yang HY, Hsu YC, Yang PC, Shih JY: EGFR-L858R mutant enhances lung adenocarcinoma cell invasive ability and promotes malignant pleural effusion formation through activation of the CXCL12-CXCR4 pathway. Sci Rep 2015, 5:13574.

43. Mitchell B, Mahalingam M: The CXCR4/CXCL12 axis in cutaneous malignancies with an emphasis on melanoma. Histol Histopathol 2014, 29(12):1539–1546.

44. Zhang J, Liu C, Mo X, Shi H, Li S: Mechanisms by which CXCR4/CXCL12 cause metastatic behavior in pancreatic cancer. Oncol Lett 2018, 15(2):1771–1776.

45. Cui K, Zhao W, Wang C, Wang A, Zhang B, Zhou W, Yu J, Sun Z, Li S: The CXCR4-CXCL12 pathway facilitates the progression of pancreatic cancer via induction of angiogenesis and lymphangiogenesis. J Surg Res 2011, 171(1):143–150.

46. Sleightholm RL, Neilsen BK, Li J, Steele MM, Singh RK, Hollingsworth MA, Oupicky D: Emerging roles of the CXCL12/CXCR4 axis in pancreatic cancer progression and therapy. Pharmacol Ther 2017, 179:158–170.

47. Feig C, Jones JO, Kraman M, Wells RJ, Deonarine A, Chan DS, Connell CM, Roberts EW, Zhao Q, Caballero OL et al: Targeting CXCL12 from FAP-expressing carcinoma-associated fibroblasts synergizes with anti-PD-L1 immunotherapy in pancreatic cancer. Proc Natl Acad Sci U S A 2013, 110(50):20212–20217.

48. Elham Azizi AJC, George Plitas, Andrew E. Cornish, Catherine Konopacki, Sandhya Prabhakaran, Juozas Nainys, Kenmin Wu, Vaidotas Kiseliovas, Manu Setty, Kristy Choi, Phuong Dao, Linas Mazutis, Alexander Y. Rudensky, Dana Pe'erdams, D. J.: Single-Cell Immune Map of Breast Carcinoma Reveals Diverse Phenotypic States Driven by the Tumor Microenvironment.

49. Gabrilovich DI: Myeloid-Derived Suppressor Cells. Cancer Immunol Res 2017, 5(1):3–8.

50. Gentles AJ, Newman AM, Liu CL, Bratman SV, Feng W, Kim D, Nair VS, Xu Y, Khuong A, Hoang CD et al: The prognostic landscape of genes and infiltrating immune cells across human cancers. Nat Med 2015, 21(8):938–945.

51. Kumar V, Donthireddy L, Marvel D, Condamine T, Wang F, Lavilla-Alonso S, Hashimoto A, Vonteddu P, Behera R, Goins MA et al: Cancer-Associated Fibroblasts Neutralize the Anti-tumor Effect of CSF1 Receptor Blockade by Inducing PMN-MDSC Infiltration of Tumors. Cancer Cell 2017, 32(5):654–668 e655.

52. Kiss M, Van Gassen S, Movahedi K, Saeys Y, Laoui D: Myeloid cell heterogeneity in cancer: not a single cell alike. Cell Immunol 2018.

53. Dobin A, Davis CA, Schlesinger F, Drenkow J, Zaleski C, Jha S, Batut P, Chaisson M, Gingeras TR: STAR: ultrafast universal RNA-seq aligner. Bioinformatics 2013, 29(1):15–21.

54. Gu Z, Eils R, Schlesner M: Complex heatmaps reveal patterns and correlations in multidimensional genomic data. Bioinformatics 2016, 32(18):2847–2849.

